# Indirect effects of parental conflict on conspecific offspring development

**DOI:** 10.1101/2021.11.18.469161

**Authors:** J.M. Coughlan

**Author notes:** Author Contributions:* JMC conceived of this project, performed all experiments, analyzed the data and wrote the paper.

## Abstract

Hybrid Seed Inviability (HSI) is a common barrier in angiosperms. Recent work suggests that the rapid evolution of HSI may, in part, be due to conflict between maternal and paternal optima for resource allocation to developing offspring (i.e. parental conflict). However, parental conflict requires that paternally-derived resource acquiring alleles impose a maternal cost. I test this requirement using three closely related species in the *Mimulus guttatus* species complex that exhibit significant HSI and differ in their inferred histories of parental conflict. I show that the presence of hybrid seeds significantly affects conspecific seed size for almost all crosses, such that conspecific seeds are smaller after developing with hybrids from fathers with a stronger history of conflict, and larger after developing with hybrids from fathers with a weaker history of conflict. This work demonstrates a cost of paternally-derived alleles, and also has implications for species fitness in secondary contact.

## Introduction

A fundamental source of conflict in viviparous organisms stems from differences between maternal and paternal optima for how many resources to allocate to developing offspring (i.e. parental conflict (Trivers 1974; Charnov 1979; Haig and Westoby 1989)). This is because in non-monogamous systems while maternity is guaranteed, fathers are not equally related to all offspring produced within a brood, and thus selection can favor the evolution of paternally-derived, resource acquiring alleles that come at a cost to either mothers directly or indirectly by influencing other developing offspring in that brood (e.g. ‘greedy alleles’; (Trivers 1974; Charnov 1979; Haig and Westoby 1989; Haig 1997; Wilkins and Haig 2001; Brandvain 2010)). Consequently, selection will then favor maternally-derived, resource repressive alleles, and a co-evolutionary arms race may subsequently evolve (Trivers 1974; Charnov 1979; Haig and Westoby 1989; Haig 1997; Wilkins and Haig 2001; Brandvain 2010). As such, parental conflict theory predicts that the severity of parental conflict is a reflection of the variance in paternity within broods (Queller 1984; Brandvain and Haig 2005; Brandvain et al. 2011; Willi 2013; Raunsgard et al. 2018). Parental conflict may be particularly important in systems where nutrients are partitioned directly, dynamically, and post-fertilization from maternal parents to developing offspring *in utero* via an intermediary tissue, such as placental or endosperm (Haig and Westoby 1989; Moore and Haig 1991; Zeh and Zeh 2000).

In seed plants, the endosperm is a nutritive tissue that is essential for proper embryo development and is analogous to the placenta in mammals. Endosperm arises via the fertilization of the central cell; a di-haploid structure within the megagametophyte, resulting in a triploid tissue that is 2 maternal genomes: 1 paternal genome. The balance of 2m:1p in the endosperm is crucial for its development, as many genes which are essential for proper endosperm development are imprinted (i.e. genes are expressed based on whether they are maternally or paternally derived), the balance of which allows development to proceed normally (Scott et al. 1998; Köhler and Weinhofer-Molisch 2010). Much research using interploidy crosses has demonstrated that an overexpression of maternally expressed or paternally expressed genes results in canonical developmental defects across multiple plant systems and endosperm developmental programs (Scott et al. 1998; Kradolfer et al. 2013; Wolff et al. 2015; Lafon-Placette and Köhler 2016; Lafon-Placette et al. 2017; Morgan et al. 2021). Strikingly, excess of maternal or paternal expression results in growth-repressive or growth-excessive phenotypes, reminiscent of predictions of parental conflict. If these endosperm defects are severe enough they can cause embryo death, and as such seed inviability is thought to be a crucial reproductive barrier between plants of different ploidy levels (often referred to as ‘Triploid Block’; (Comai 2005; Köhler et al. 2010, 2021; Sutherland and Galloway 2017; Morgan et al. 2021)).

Yet, Hybrid Seed Inviability (HSI) is also common in diploid plant systems and largely results from parent-of-origin specific growth defects in the endosperm (Brandvain and Haig 2005; Lowry et al. 2008; Briscoe Runquist et al. 2014; Rebernig et al. 2015; Garner et al. 2016; Lafon-Placette and Köhler 2016; Oneal et al. 2016; Lafon-Placette et al. 2017, 2018; Roth et al. 2018*b*, 2019; Coughlan et al. 2020*b*; Sandstedt et al. 2020; Gustafsson et al. 2021; İltaş et al. 2021). These patterns, while strikingly similar to the defects exemplified in interploidy crosses (Lafon-Placette and Köhler 2016; Lafon-Placette et al. 2017; Städler et al. 2021), cannot be explained by genome-wide imbalances of maternal:paternal gene expression in the endosperm, and must involve the evolution of paternal-excessive and maternal-repressive alleles. The observation that interspecific diploid crosses often mirror inter-ploidy crosses has sparked a conceptual framework to categorize diploid taxa according to the extent of maternal-repression and paternal-excess that they exhibit when crossed to other diploids, referred to as their Endosperm Balance Number (EBN;(Johnston et al. 1980; Katsiotis et al. 1995; Carputo et al. 1999; Johnston and Hanneman 1999; Lafon-Placette and Köhler 2016; Lafon-Placette et al. 2018; Städler et al. 2021)), or genome strength (Brandvain and Haig 2018). Differences between taxa in EBNs are thought to reflect different histories of parental conflict (Lafon-Placette et al. 2018; Raunsgard et al. 2018; Coughlan et al. 2020*b*; Städler et al. 2021).

Although its role in underlying HSI is garnering much current support (Lafon-Placette et al. 2018; Coughlan et al. 2020*b*; İltaş et al. 2021), parental conflict may also play a secondary role in speciation in the context of secondary contact. Hybridization between closely related species that vary in EBNs can not only result in the loss of gametes (i.e. reproductive interference), but when hybrids are formed in a brood that also contains conspecific offspring, differences in the competitive ability between hybrid and conspecific siblings borne through differences in paternally-derived resource acquiring alleles may affect conspecific size, and consequently offspring fitness. In seed plants, seed size is often a proxy for various fitness components (such as the probability of germination, seedling size, and the number of flowers produced (Stanton 1984; Krannitz et al. 1991; Simons and Johnston 2000; Gómez 2004)). Thus, competition for limited maternal resources between hybrid and intraspecific siblings may have substantial implications for intraspecific fitness in secondary contact zones.

Here I use a model organism for ecology, evolution, and genetics; the *Mimulus guttatus* species complex to address if conspecific seed size varies when seeds are grown only with other conspecific siblings versus hybrid siblings from sires with different EBNs. Previously, I have shown that *M. guttatus* and a closely related species, *M. decorus,* are reproductively isolated by HSI, and patterns of HSI support a role for parental conflict (Coughlan et al. 2020*b*). Namely, hybrid seeds exhibit parent-of-origin specific growth defects (maternal-repressive and paternal-excess phenotypes) that are associated with parent-of-origin specific endosperm defects (Coughlan et al. 2020*b*). *Mimulus decorus* comprises two distinct diploid lineages; one that exhibits substantially lower EBN than *M. guttatus* and one that exhibits a higher EBN than *M. guttatus*, despite a relatively recent divergence time (roughly 230kya; (Coughlan et al. 2020*b*)). Here, I leverage this diversity of EBNs in this group to assess indirect growth effects in conspecific seeds when these seeds are grown alongside hybrid siblings that vary in their father’s EBN. This work is one of the first to provide an explicit test of the cost of paternally derived, resource acquiring alleles; a central prediction of parental conflict. Moreover, the results of this experiment have implications for secondary contact between species that differ in EBN.

## Materials & Methods

### Plant materials and crosses

I grew between 6-16 replicate lines of a single genotype for each of *M. guttatus* (*IM62*), Southern *M. decorus* (*OD11*), and Northern *M. decorus* (*HWY15D*). Previous work using population genomics and experimental crosses suggests that Northern *M. decorus* produces relatively weak dams and sires (i.e. has a low EBN), and is predicted to have experienced the weakest history of parental conflict, while Southern *M. decorus* produces relatively strong dams and sires (i.e. a high EBN), and is predicted to have experienced the strongest history of parental conflict (Coughlan et al. 2020*b*). *Mimulus guttatus* exhibits an EBN intermediate to the two genetic lineages of *M. decorus* (i.e. stronger than Northern *M. decorus,* but weaker than Southern *M. decorus*; *(Coughlan et al. 2020b)*). I first cold and dark stratified seeds in water for one week, then dispersed them on moist Fafard 4P soil (SunGro Horticultural Inc) in the University of North Carolina at Chapel Hill greenhouses. I transferred early germinants to individual 2 1/2” pots and grew plants in warm, long day conditions (16hours of light/ 20C).

I emasculated individual flowers at least one day before flowers would have naturally opened and well before natural pollen dehiscence. All species used in this experiment have a highly outcrossing morphology and very rarely (if ever) set autogamous selfed seed in the greenhouse (Coughlan and Willis 2019; Coughlan et al. 2020*b*). In accordance with this observation, no unpollinated stigma bore a fruit during the course of this experiment. For each species, stigmas were pollinated with either pure self pollen or a mixture of self pollen and pollen from another species. For both pollination types, anthers were manually dehisced on a glass slide, then I manually transferred pollen to an open stigma using fine forceps. For the mixed pollination treatment, anthers were manually dehisced until a roughly equal quantity of pollen from each species was available, then pollen samples were thoroughly mixed with fine forceps. The exact ratio of pollen from each parent is unlikely to have been equal for every cross, and indeed fertilization rates between intra- and inter-specific fathers did significantly differ for may fruits, which may reflect departures from a 1:1 ratio of pollen genotypes or differences in competitive ability of pollen genotypes (see Table S2). Nonetheless, the aim of this work is to assess if the presence of hybrid siblings influences intraspecific seed size, and as such an exact 1:1 ratio of paternal genotypes in the pollen is unnecessary. I collected fully ripened fruits shortly before natural dehiscence. All fruits were dried at ambient temperatures for at least one week before seeds were counted, categorized, and measured. In total, an average of 26.5 fruits (range: 20-31 fruits) were quantified for each species and cross type combination (see Figure 1 for overview of experimental design).

**Figure 1:**
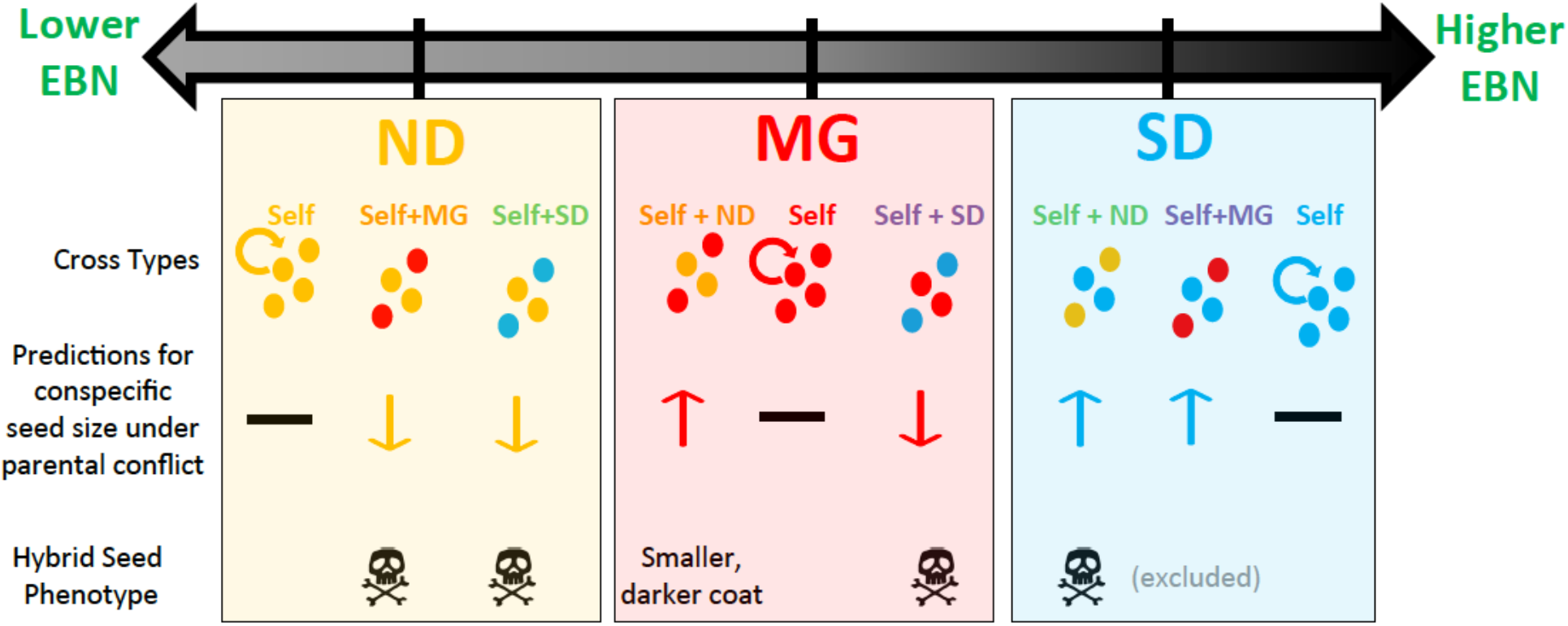
Overview of the experimental design. This experiment leveraged natural variation in EBN across three species (ND= Northern *M. decorus*, MG= *M. guttatus*, SD= Southern *M. decorus*; outlined in (Coughlan et al. 2020*b*)). To compare conspecific seed size when conspecific seeds developed along full siblings versus half siblings whose sires vary in EBN, nine types of crosses were completed: each species was self fertilized, or fertilized with a mixture of self-pollen and pollen from another species. After excluding putative hybrid seeds based on diagnostic phenotypes (hybrids having inviable or much smaller and darker seeds; see Figure S1 for confirmation), all conspecific seeds for each cross type were measured for total area. Prediction based on parental conflict theory are outlined: conspecific seeds developing with hybrid siblings sired by a higher EBN father should be smaller (exemplified by both mixed crosses involving Northern *M. decorus* as the dam), while conspecific seeds developing alongside hybrid siblings sired by a lower EBN father should be larger (exemplified by both mixed crosses involving Southern *M. decorus* as the dam).

### Seed Quantification

For each cross, all fertilized seeds were counted and quantified for viability based on morphology, with shriveled, disc-like, or concave seeds scored as inviable. Morphology has previously been shown to be a good indicator of viability in *Mimulus* (Garner et al. 2016; Coughlan et al. 2020*b*). In four of the six mixed pollen treatments, hybrids are almost always inviable (<<<0.1% viability. In contrast, selfed crosses show generally low rates of inviability (~95% viable); (Coughlan et al. 2020*b*)). We therefore assumed that any viable seed in these four mixed pollination crosses were likely to be a product of self fertilization. In two of the six mixed-pollen crosses, hybrid seeds are generally viable, though noticeably smaller and produce much darker seed coats than selfed seed (*M. guttatus* mothers x Northern *M. decorus* pollen donors and Southern *M. decorus* mothers x *M. guttatus* pollen donors). In the case of Southern *M. decorus,* crosses in which pollen was mixed with *M. guttatus* yielded a generally high level of fruit failure, and thus this cross type was excluded from subsequent analysis. In the case of crosses involving *M. guttatus* mothers with a mixture of *M. guttatus* and Northern *M. decorus* pollen, a subset of putative self-fertilized and putative hybrid seeds were grown to confirm that designations based on seed phenotype accurately indicated paternal genotype. All seeds which were presumed to be self fertilized were indeed conspecific seeds, and only 2/86 presumed hybrids were likely conspecific seeds (~2.3% error rate; Figure S1). I then measured the area of all viable, putatively non-hybrid seeds using ImageJ (Schneider et al. 2012) for the remaining 8 cross type/maternal parent combinations. In total, this resulted in an average of 29, 55, and 128 seeds measured per fruit, for crosses involving Southern *M. decorus*, *M. guttatus*, and Northern *M. decorus* as the maternal donor, respectively. In total, I measured 16,199 seeds across 215 fruits for all 8 experimental cross types.

### Analyses

To assess if the presence of hybrid siblings affected the size of intraspecific seeds, I used a linear mixed model with log(seed area) as the response variable and cross type (e.g. pure self-fertilized, mixed pollen with species 1 and mixed pollen with species 2 as the levels), and the total number of seeds per fruit as fixed effects. As the same plant was used for multiple crosses, and multiple seeds were measured from a single fruit, I also included maternal replicate and fruit replicate as random effects. The significance of the fixed effects were assessed by a Type III Wald ***χ*^2^** test using the *lme4* and *car* packages in R (Bates et al. 2012; Fox and Weisberg 2018). Significance between cross types were assessed using the *emmeans* package in R (Russell 2019). These analyses were performed for each maternal genotype separately.

## Results

### The presence of hybrid seeds affects intraspecific seed size

The presence of hybrid seeds significantly affected conspecific seed size in almost all crosses (Figure 2, Table S1). The direction of these effects depended on the type of hybrids that conspecific seeds developed alongside (Figure 2). For *M. guttatus*, conspecific seeds were smaller when they developed along hybrids sired by a higher EBN pollen donor (i.e. Southern *M. decorus*), and larger when they developed alongside hybrid siblings sired by a lower EBN pollen donor (i.e. Northern *M. decorus*; Figure 2). Similarly, for Southern *M. decorus*, conspecific seeds were larger when they developed alongside hybrid siblings sired by a lower EBN pollen donor (i.e. Northern *M. decorus*; Figure 2). For Northern *M. decorus*, conspecific seeds were significantly smaller when developing alongside hybrid siblings sired by the highest EBN pollen donor in this experiment (i.e. Southern *M. decorus*; Figure 2), although there is no significant difference in intraspecific seed size between fruits containing only pure intraspecific selfs and fruits containing both intraspecific selfs and hybrid siblings sired by *M. guttatus* (which has a higher EBN than Northern *M. decorus*, but a lower EBN that Southern *M. decorus*). Overall, these effects are relatively small, for example intraspecific *M. guttatus* seeds are ~5% larger when developing alongside hybrids from a lower EBN father and ~7% smaller when developing alongside hybrids from a higher EBN father (see Figure 2 for all estimated effect sizes based on estimated marginal means).

**Figure 2:**
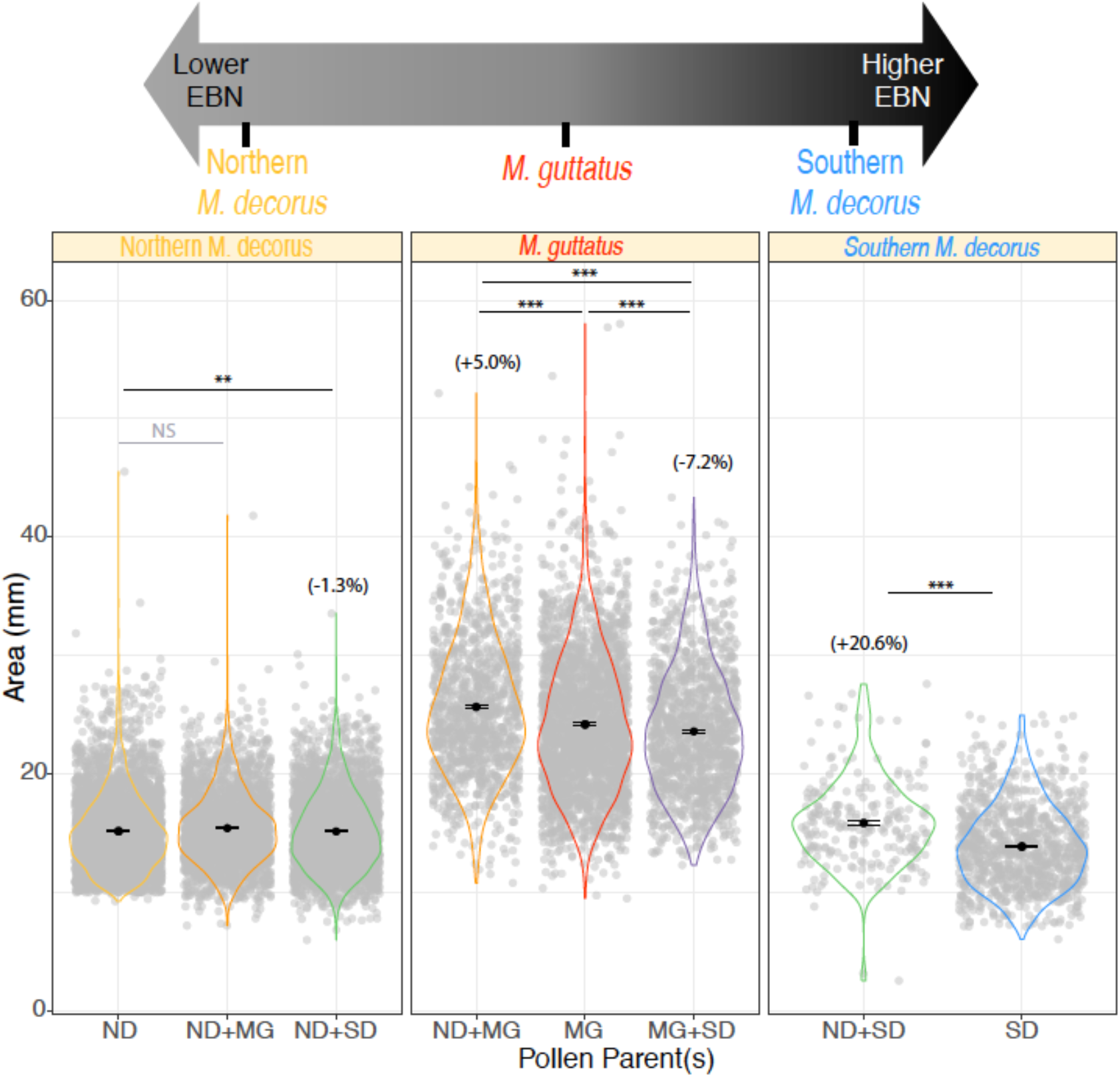
Intraspecific seed size differs between developmental contexts. For each species, either pure conspecific (self) pollen or a mixture of con- and interspecific pollen was applied to receptive stigmas, then the resultant conspecific seeds were measured for total area (in mm). Each panel depicts seeds produced from each genotype as the maternal parent, while colors of the violins indicate the pollen donor(s) (note: while only conspecific seeds were measured, for mixed pollination crosses I use an intermediate color between the two pollen donors for contrast to pure conspecific crosses). Filled, solid black points represent the means and standard errors, while translucent, grey points are all raw data. For crosses that show a significant difference in conspecific seed size, the effect size relative to conspecific seeds that developed alongside full siblings as estimated from the *emmeans* package in R (Russell 2019) are shown in brackets. Significant differences were determined using a linear mixed effect model (See Table S1 for output). Note that for Southern *M. decorus* fruits generally failed when fertilized with a combination of self-pollen and pollen from *M. guttatus*, so this cross was excluded from further analysis.

## Discussion

Here I show that conspecific seed size is influenced by the presence of hybrid siblings during development. These results are consistent with the idea that paternally-derived, resource acquiring alleles are costly to less resource-competitive siblings developing in the same brood (and in turn, to maternal parents). Although these maternal costs are a central prediction of parental conflict theory, very few studies have empirically demonstrated them. To my knowledge, the only other study to show these costs utilized variation between maize accessions for paternally-derived resource acquiring alleles to show that seed size differed when half-siblings whose fathers differed in their competitive ability developed alongside one another relative to when the same genotypes developed alongside full siblings (Cailleau et al. 2018). Here, I leverage proposed differences in EBN between recently diverged taxa to illustrate a similar phenomenon.

While a growing body of evidence has highlighted the role of parental conflict in the origins of reproductive isolation (namely HSI, and early-onset hybrid inviability in mammalian systems; (Vrana et al. 1998, 2000; Brekke and Good 2014; Brekke et al. 2016, 2021; Oneal et al. 2016; Lafon-Placette et al. 2018; Roth et al. 2018*a*; Coughlan et al. 2020*b*; Sandstedt et al. 2020; Arévalo et al. 2021)), the work presented here highlights a secondary role of parental conflict in speciation: hybridization between species that vary in their histories of parental conflict may result in indirect effects to intraspecific offspring development, in the event that intraspecific offspring develop alongside hybrids. In this system, *M. guttatus* co-occurs with both Northern and Southern *M. decorus* for large portions of their range (JMC personal obs; (Coughlan et al. 2020*a*)), and are thought to routinely hybridize and introgress (JMC unpublished data; (Puzey et al. 2017)). Although seed size differences have been shown to influence several components of fitness in other systems (germination probability, seedling survivorship, flower production; (Stanton 1984; Simons and Johnston 2000; Gómez 2004)), these effects are likely to be context specific (for example, based on the competitive environment; (Stanton 1984)), and substantial fieldwork and further experimentation is required to quantify fitness effects in this system. However, one intriguing finding of this work is that the consequences of hybridization for intraspecific seed development are not always costs. If hybridization with fathers with lower EBNs consistently results in slightly larger conspecific seeds, this hybridization might actually result in a fitness benefit to individual seeds. However, this potential benefit is likely a very small one, and moreover hybridization likely still results in an overall cost to maternal parents and a loss of inclusive fitness to individual seeds, given the loss of gametes to inviable hybrid offspring. Depending on the rates of hybridization and the competitive environment in which seeds (or in the case of mammals, young offspring) find themselves, this work may have significant implications for the dynamics of hybridization and coexistence of species that vary in their histories of conflict and find themselves in secondary contact.

## Data Availability

All raw data are available under the Dryad submission https://doi.org/10.5061/dryad.m905qfv2d. Summaries of all crosses are included in Supplementary Table 2 of this manuscript.

## Acknowledgements

Thanks to the Willis, Sweigart, Sobel, Franks and Matute labs for providing helpful feedback on this manuscript. I would also like to graciously thank Polly Campbell and Sarah Gardner for a particularly creative seminar question that greatly inspired my thought process in designing this experiment. JMC was supported by an NSF DEB Dimensions of Biodiversity grant awarded to Daniel Matute (DEB-DOB-1737752).

## Supplemental Figures

**Figure S1:**
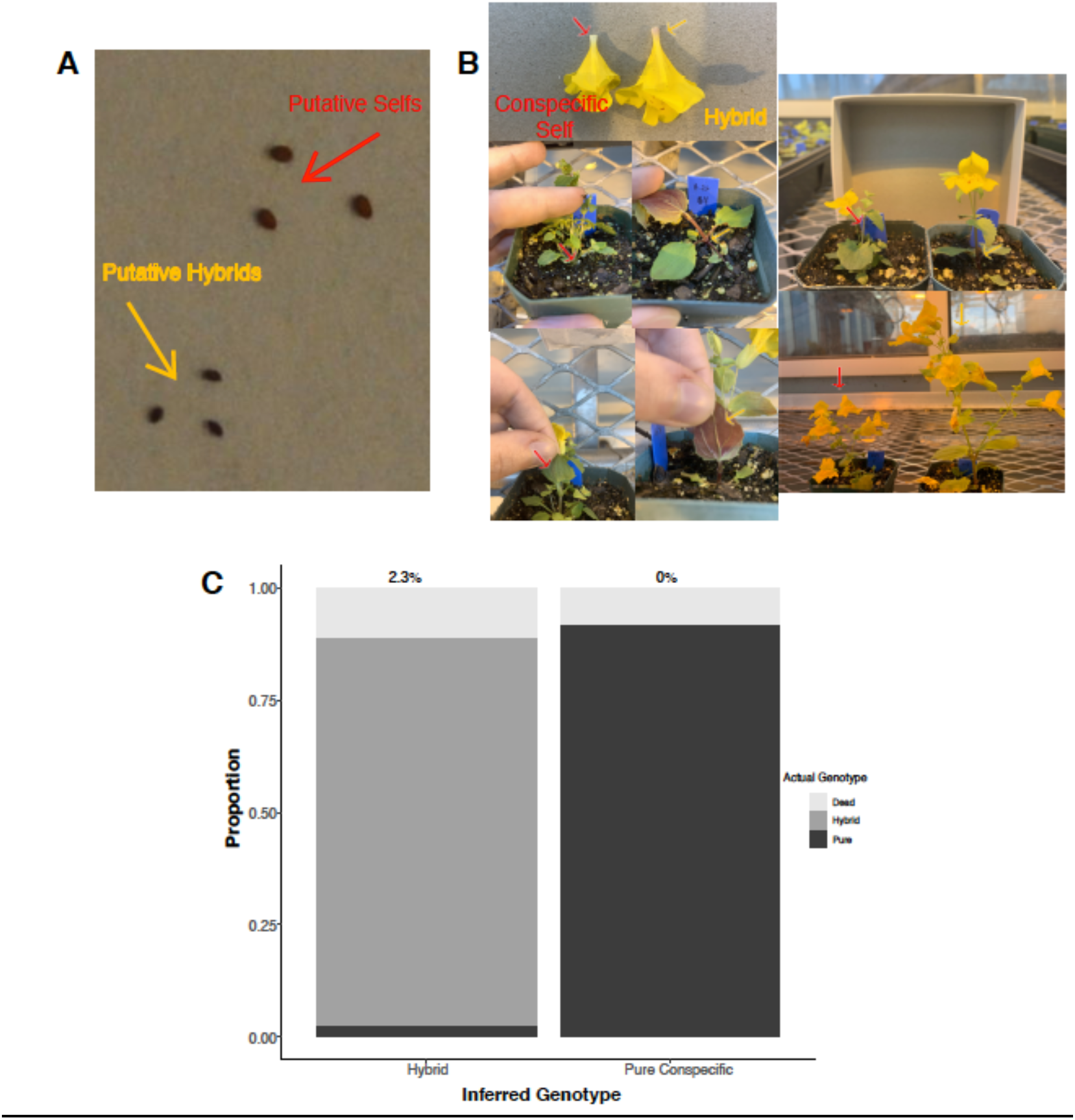
Confirmation of paternal genotypes for mixed pollination crosses involving *M. guttatus* as the dam and mixed pollen from *M. guttatus* and Northern *M. decorus*. (A) Putative hybrids and putative self-fertilized seeds exhibit distinct seed characteristics; hybrids are smaller and have much darker seed coats. To confirm paternal genotype, a subset of putative selfs and putative hybrids were grown. Plants differ in a number of diagnostic characters (selfs on the left, hybrids on the right), including: presence anthocyanin pigmentation in the base of the corolla in hybrids (top left), the presence of stolons in hybrids (middle left), the presence of anthocyanin pigmentation on the underside of the leaf in hybrids (bottom left), stems with anthocyanin pigmentation in hybrids (top right), and overall size; both of flowers and total plant, wherein hybrids are larger (bottom right). (C) Based on this set of characters, there were no hybrids that were designated as selfed seed and only ~2% of seeds designated as hybrids were in fact conspecific seeds.

## Supplemental Tables

**Table S1:**
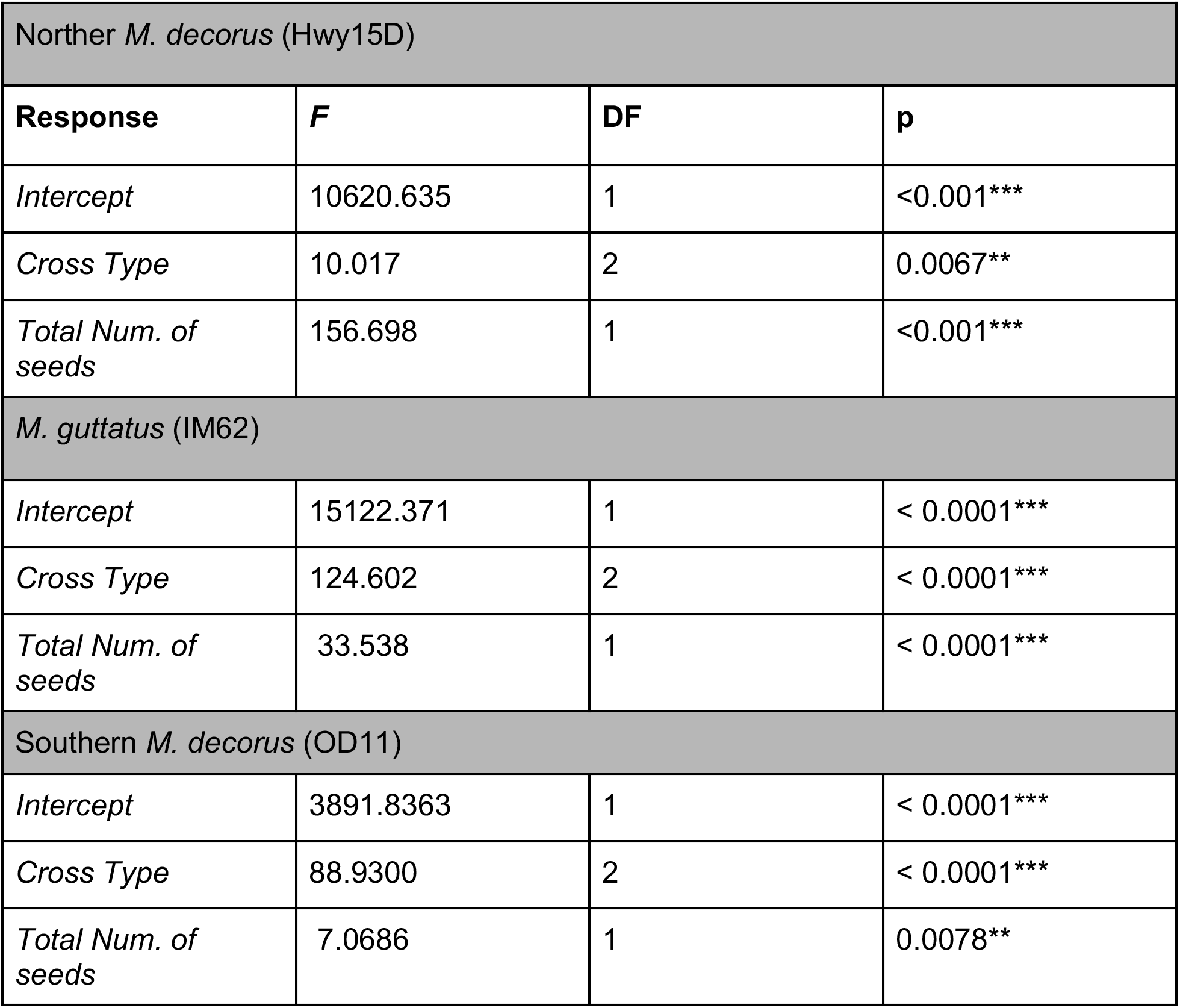
LMER outputs for each species as maternal parent. In all cases, log(seed area) was the response variable, with Cross Type (pure-self pollinated, mixed pollen with species 1, mixed pollen with species 2 as the levels), and the total number of seeds per fruit as fixed effects. Maternal plant replicate and fruit replicate were included as random effects. Significance of fixed effects was determined using Type III Wald ***χ*^2^** test using the lme4 and car packages in R (Bates et al. 2012; Fox and Weisberg 2018).

| Norther *M. decorus* (Hwy15D) |  |  |  |
| --- | --- | --- | --- |
| Response | F | DF | p |
| Intercept | 10620.635 | 1 | <0.001*** |
| Cross Type | 10.017 | 2 | 0.0067** |
| Total Num. of seeds | 156.698 | 1 | <0.001*** |
| *M. guttatus* (IM62) |  |  |  |
| Intercept | 15122.371 | 1 | < 0.0001*** |
| Cross Type | 124.602 | 2 | < 0.0001*** |
| Total Num. of seeds | 33.538 | 1 | < 0.0001*** |
| Southern *M. decorus* (OD11) |  |  |  |
| Intercept | 3891.8363 | 1 | < 0.0001*** |
| Cross Type | 88.9300 | 2 | < 0.0001*** |
| Total Num. of seeds | 7.0686 | 1 | 0.0078** |

**Table S2:** Number of seeds produced, proportion inviable, and average seed size per fruit for all crosses. **Mat**= Maternal genotype; **Mat.Rep**= individual replicate of that genotype; **Pat**= Paternal donor(s); **CR**= Cross Replicate (fruit replicate for each Maternal/Paternal combination); **Viable**= Number of viable seeds; **Inviable**= Number of inviable seeds; **Total**= total number of seeds produced per fruit; **Prop**= proportion of inviable seeds; **AA**= average seed area (mm), **Sig**= is there a significant deviation from 1:1 pollination of the two paternal genotypes (SIG= significant deviation, NS= Not-significant, NA= not applicable (i.e. only one paternal donor))).

| Mat | Mat.Rep | Pat | CR | Viable | Inviabile | Total | Prop | AA | Sig? |
| --- | --- | --- | --- | --- | --- | --- | --- | --- | --- |
| IM62 | 4 | OD+IM | 1 | 20 | 8 | 28 | 0.2857 | 21.6246 | NS |
| IM62 | 14 | OD+IM | 2 | 90 | 0 | 90 | 0.0000 | 18.5862 | SIG |
| IM62 | 0 | OD+IM | 3 | 3 | 7 | 10 | 0.7000 | 31.9987 | NS |
| IM62 | 12 | OD+IM | 4 | 15 | 8 | 23 | 0.3478 | 29.3106 | NS |
| IM62 | 0 | OD+IM | 5 | 117 | 15 | 132 | 0.1136 | 21.2291 | SIG |
| IM62 | 0 | OD+IM | 6 | 62 | 9 | 71 | 0.1268 | 25.6564 | SIG |
| IM62 | 0 | OD+IM | 7 | 48 | 62 | 110 | 0.5636 | 22.2014 | NS |
| IM62 | 13 | OD+IM | 8 | 28 | 15 | 43 | 0.3488 | 24.1408 | NS |
| IM62 | 6 | OD+IM | 9 | 28 | 25 | 53 | 0.4717 | 23.7778 | NS |
| IM62 | 12 | OD+IM | 10 | 39 | 47 | 86 | 0.5465 | 24.0505 | NS |
| IM62 | 5 | OD+IM | 11 | 34 | 46 | 80 | 0.5750 | 25.1875 | NS |
| IM62 | 14 | OD+IM | 12 | 18 | 53 | 71 | 0.7465 | 21.0091 | SIG |
| IM62 | 13 | OD+IM | 13 | 38 | 23 | 61 | 0.3770 | 26.2518 | NS |
| IM62 | 13 | OD+IM | 14 | 7 | 37 | 44 | 0.8409 | 24.7432 | SIG |
| IM62 | 2 | OD+IM | 15 | 14 | 23 | 37 | 0.6216 | 24.0467 | NS |
| IM62 | 0 | OD+IM | 16 | 11 | 7 | 18 | 0.3889 | 20.1029 | NS |
| IM62 | 3 | OD+IM | 17 | 6 | 10 | 16 | 0.6250 | 29.2899 | NS |
| IM62 | 0 | OD+IM | 18 | 24 | 6 | 30 | 0.2000 | 24.5210 | NS |
| IM62 | 7 | OD+IM | 19 | 47 | 14 | 61 | 0.2295 | 23.1699 | SIG |
| IM62 | 15 | OD+IM | 20 | 26 | 11 | 37 | 0.2973 | 22.3076 | NS |
| IM62 | 6 | OD+IM | 21 | 41 | 23 | 64 | 0.3594 | 25.3321 | NS |
| IM62 | 6 | OD+IM | 22 | 6 | 0 | 6 | 0.0000 | 25.4753 | NS |
| IM62 | 9 | OD+IM | 23 | 112 | 8 | 120 | 0.0667 | 23.5290 | SIG |
| IM62 | 8 | OD+IM | 24 | 11 | 12 | 23 | 0.5217 | 23.7891 | NS |
| IM62 | 4 | OD+IM | 25 | 5 | 25 | 30 | 0.8333 | 27.2043 | NS |
| IM62 | 0 | OD+IM | 26 | 44 | 11 | 55 | 0.2000 | 24.6987 | SIG |
| IM62 | 0 | OD+IM | 27 | 92 | 10 | 102 | 0.0980 | 25.9432 | SIG |
| IM62 | 0 | OD+IM | 28 | 48 | 20 | 68 | 0.2941 | 22.2567 | NS |
| IM62 | 0 | OD+IM | 29 | 38 | 19 | 57 | 0.3333 | 29.3798 | NS |
| IM62 | 0 | OD+IM | 30 | 41 | 36 | 77 | 0.4675 | 26.0528 | NS |
| HWY15D | 6 | IM62+HWY | 1 | 182 | 157 | 339 | 0.4631 | 13.7714 | NS |
| HWY15D | 4 | IM62+HWY | 2 | 42 | 96 | 138 | 0.6957 | 16.8653 | SIG |
| HWY15D | 6 | IM62+HWY | 3 | 91 | 54 | 145 | 0.3724 | 17.2701 | NS |
| HWY15D | 4 | IM62+HWY | 4 | 139 | 144 | 283 | 0.5088 | 15.3169 | NS |
| HWY15D | 4 | IM62+HWY | 5 | 84 | 160 | 244 | 0.6557 | 15.4467 | SIG |
| HWY15D | 6 | IM62+HWY | 6 | 58 | 61 | 119 | 0.5126 | 17.5574 | NS |
| HWY15D | 6 | IM62+HWY | 7 | 65 | 99 | 164 | 0.6037 | 15.2325 | NS |
| HWY15D | 6 | IM62+HWY | 8 | 72 | 61 | 133 | 0.4586 | 16.7600 | NS |
| HWY15D | 6 | IM62+HWY | 9 | 101 | 110 | 211 | 0.5213 | 16.2864 | NS |
| HWY15D | 4 | IM62+HWY | 10 | 63 | 134 | 197 | 0.6802 | 17.3873 | SIG |
| HWY15D | 4 | IM62+HWY | 11 | 44 | 122 | 166 | 0.7349 | 17.8447 | SIG |
| HWY15D | 4 | IM62+HWY | 12 | 30 | 79 | 109 | 0.7248 | 18.8440 | SIG |
| HWY15D | 6 | IM62+HWY | 13 | 22 | 23 | 45 | 0.5111 | 15.9900 | NS |
| HWY15D | 6 | IM62+HWY | 14 | 141 | 105 | 246 | 0.4268 | 14.9731 | NS |
| HWY15D | 6 | IM62+HWY | 15 | 58 | 36 | 94 | 0.3830 | 16.2463 | NS |
| HWY15D | 7 | IM62+HWY | 16 | 149 | 94 | 243 | 0.3868 | 17.2735 | NS |
| HWY15D | 5 | IM62+HWY | 17 | 96 | 135 | 231 | 0.5844 | 13.7065 | NS |
| HWY15D | 5 | IM62+HWY | 18 | 61 | 108 | 169 | 0.6391 | 14.1027 | NS |
| HWY15D | 7 | IM62+HWY | 19 | 80 | 62 | 142 | 0.4366 | 17.4737 | NS |
| HWY15D | 7 | IM62+HWY | 20 | 46 | 32 | 78 | 0.4103 | 16.4966 | NS |
| HWY15D | 7 | IM62+HWY | 21 | 124 | 72 | 196 | 0.3673 | 16.3268 | NS |
| HWY15D | 7 | IM62+HWY | 22 | 15 | 57 | 72 | 0.7917 | 16.2255 | SIG |
| HWY15D | 4 | IM62+HWY | 23 | 82 | 163 | 245 | 0.6653 | 15.9680 | SIG |
| HWY15D | 6 | IM62+HWY | 24 | 132 | 52 | 184 | 0.2826 | 13.3957 | SIG |
| HWY15D | 6 | IM62+HWY | 25 | 214 | 137 | 351 | 0.3903 | 13.7087 | SIG |
| HWY15D | 4 | IM62+HWY | 26 | 37 | 141 | 178 | 0.7921 | 18.2464 | SIG |
| HWY15D | 4 | IM62+HWY | 27 | 10 | 12 | 22 | 0.5455 | 21.0723 | NS |
| HWY15D | 6 | IM62+HWY | 28 | 36 | 43 | 79 | 0.5443 | 13.8636 | NS |
| HWY15D | 4 | IM62+HWY | 29 | 49 | 40 | 89 | 0.4494 | 16.5796 | NS |
| HWY15D | 6 | IM62+HWY | 30 | 99 | 98 | 197 | 0.4975 | 13.4846 | NS |
| HWY15D | 6 | OD+HWY15 | 1 | 166 | 65 | 231 | 0.2814 | 14.7041 | SIG |
| HWY15D | 6 | OD+HWY15 | 2 | 229 | 145 | 374 | 0.3877 | 14.6194 | SIG |
| HWY15D | 6 | OD+HWY15 | 3 | 276 | 152 | 428 | 0.3551 | NA | SIG |
| HWY15D | 6 | OD+HWY15 | 4 | 206 | 110 | 316 | 0.3481 | 14.5547 | SIG |
| HWY15D | 4 | OD+HWY15 | 5 | 30 | 90 | 120 | 0.7500 | 16.4631 | SIG |
| HWY15D | 4 | OD+HWY15 | 6 | 92 | 88 | 180 | 0.4889 | 17.2475 | NS |
| HWY15D | 6 | OD+HWY15 | 7 | 48 | 32 | 80 | 0.4000 | 17.5165 | NS |
| HWY15D | 6 | OD+HWY15 | 8 | 183 | 78 | 261 | 0.2989 | 15.0614 | SIG |
| HWY15D | 4 | OD+HWY15 | 9 | 19 | 19 | 38 | 0.5000 | 20.1781 | NS |
| HWY15D | 4 | OD+HWY15 | 10 | 217 | 161 | 378 | 0.4259 | 14.3388 | NS |
| HWY15D | 4 | OD+HWY15 | 11 | 231 | 162 | 393 | 0.4122 | 13.8283 | NS |
| HWY15D | 4 | OD+HWY15 | 12 | 121 | 71 | 192 | 0.3698 | 15.8368 | NS |
| HWY15D | 6 | OD+HWY15 | 13 | 64 | 57 | 121 | 0.4711 | 14.9187 | NS |
| HWY15D | 6 | OD+HWY15 | 14 | 29 | 32 | 61 | 0.5246 | 15.5215 | NS |
| HWY15D | 6 | OD+HWY15 | 15 | 37 | 12 | 49 | 0.2449 | 17.8154 | NS |
| HWY15D | 6 | OD+HWY15 | 16 | 26 | 57 | 83 | 0.6867 | 14.6550 | NS |
| HWY15D | 4 | OD+HWY15 | 17 | 265 | 96 | 361 | 0.2659 | 15.5190 | SIG |
| HWY15D | 4 | OD+HWY15 | 18 | 313 | 132 | 445 | 0.2966 | 14.8546 | SIG |
| HWY15D | 6 | OD+HWY15 | 19 | 129 | 107 | 236 | 0.4534 | 16.2029 | NS |
| HWY15D | 5 | OD+HWY15 | 20 | 10 | 39 | 49 | 0.7959 | 17.1143 | SIG |
| HWY15D | 5 | OD+HWY15 | 21 | 126 | 166 | 292 | 0.5685 | 14.4543 | NS |
| HWY15D | 5 | OD+HWY15 | 22 | 64 | 134 | 198 | 0.6768 | 13.1848 | SIG |
| HWY15D | 6 | OD+HWY15 | 23 | 126 | 96 | 222 | 0.4324 | 15.9587 | NS |
| HWY15D | 6 | OD+HWY15 | 24 | 88 | 72 | 160 | 0.4500 | 19.0036 | NS |
| HWY15D | 4 | OD+HWY15 | 25 | 53 | 72 | 125 | 0.5760 | 15.0604 | NS |
| OD11 | 3 | HWY15+OD | 1 | 64 | 104 | 168 | 0.6190 | 14.1142 | NS |
| OD11 | 3 | HWY15+OD | 2 | 6 | 32 | 38 | 0.8421 | 15.3353 | SIG |
| OD11 | 3 | HWY15+OD | 3 | 16 | 110 | 126 | 0.8730 | 15.9940 | SIG |
| OD11 | 5 | HWY15+OD | 4 | 12 | 95 | 107 | 0.8879 | 15.9141 | SIG |
| OD11 | 5 | HWY15+OD | 5 | 6 | 121 | 127 | 0.9528 | 14.3608 | SIG |
| OD11 | 5 | HWY15+OD | 6 | 8 | 35 | 43 | 0.8140 | 14.8058 | SIG |
| OD11 | 5 | HWY15+OD | 7 | 8 | 97 | 105 | 0.9238 | 14.8111 | SIG |
| OD11 | 5 | HWY15+OD | 8 | 12 | 154 | 166 | 0.9277 | 15.4242 | SIG |
| OD11 | 5 | HWY15+OD | 9 | 4 | 67 | 71 | 0.9437 | 15.9485 | SIG |
| OD11 | 3 | HWY15+OD | 10 | 23 | 137 | 160 | 0.8563 | 17.6589 | SIG |
| OD11 | 3 | HWY15+OD | 11 | 13 | 36 | 49 | 0.7347 | 19.9525 | NS |
| OD11 | 3 | HWY15+OD | 12 | 2 | 19 | 21 | 0.9048 | 14.7240 | SIG |
| OD11 | 5 | HWY15+OD | 13 | 20 | 74 | 94 | 0.7872 | 15.2884 | SIG |
| OD11 | 5 | HWY15+OD | 14 | 8 | 61 | 69 | 0.8841 | 17.9506 | SIG |
| OD11 | 3 | HWY15+OD | 15 | 7 | 20 | 27 | 0.7407 | 15.7819 | NS |
| OD11 | 5 | HWY15+OD | 16 | 7 | 53 | 60 | 0.8833 | 15.3839 | SIG |
| OD11 | 3 | HWY15+OD | 17 | 10 | 74 | 84 | 0.8810 | 17.4214 | SIG |
| OD11 | 5 | HWY15+OD | 18 | 2 | 54 | 56 | 0.9643 | 18.0635 | SIG |
| OD11 | 5 | HWY15+OD | 19 | 2 | 51 | 53 | 0.9623 | 15.7985 | SIG |
| OD11 | 5 | HWY15+OD | 20 | 6 | 26 | 32 | 0.8125 | 18.4395 | NS |
| OD11 | 3 | HWY15+OD | 21 | 4 | 11 | 36 | 0.3056 | 19.8088 | NS |
| OD11 | 3 | HWY15+OD | 22 | 0 | 11 | 11 | 1.0000 |  | NS |
| IM62 | 0 | IM62+HWY | 1 | 67 | 108 | 175 | 0.6171 | 25.0784 | NS |
| IM62 | 7 | IM62+HWY | 2 | 34 | 26 | 60 | 0.4333 | 15.3029 | NS |
| IM62 | 9 | IM62+HWY | 3 | 64 | 41 | 105 | 0.3905 | 23.5853 | NS |
| IM62 | 15 | IM62+HWY | 4 | 117 | 73 | 190 | 0.3842 | 22.9275 | NS |
| IM62 | 8 | IM62+HWY | 5 | 42 | 51 | 93 | 0.5484 | 21.2626 | NS |
| IM62 | 5 | IM62+HWY | 6 | 5 | 11 | 16 | 0.6875 | 23.9093 | NS |
| IM62 | 12 | IM62+HWY | 7 | 56 | 35 | 91 | 0.3846 | 23.5435 | NS |
| IM62 | 9 | IM62+HWY | 8 | 7 | 18 | 25 | 0.7200 | 28.8460 | NS |
| IM62 | 15 | IM62+HWY | 9 | 6 | 19 | 25 | 0.7600 | 27.4792 | NS |
| IM62 | 5 | IM62+HWY | 10 | 33 | 35 | 68 | 0.5147 | 28.8596 | NS |
| IM62 | 5 | IM62+HWY | 11 | 43 | 121 | 164 | 0.7378 | 28.8578 | SIG |
| IM62 | 12 | IM62+HWY | 12 | 67 | 157 | 224 | 0.7009 | 21.8156 | SIG |
| IM62 | 11 | IM62+HWY | 13 | 36 | 34 | 70 | 0.4857 | 25.0363 | NS |
| IM62 | 12 | IM62+HWY | 14 | 41 | 61 | 102 | 0.5980 | 30.0482 | NS |
| IM62 | 0 | IM62+HWY | 15 | 8 | 22 | 30 | 0.7333 | 22.3832 | NS |
| IM62 | 0 | IM62+HWY | 16 | 27 | 79 | 106 | 0.7453 | 26.9391 | SIG |
| IM62 | 0 | IM62+HWY | 17 | 18 | 17 | 35 | 0.4857 | 34.5770 | NS |
| IM62 | 0 | IM62+HWY | 18 | 34 | 67 | 101 | 0.6634 | 32.6350 | NS |
| IM62 | 0 | IM62+HWY | 19 | 42 | 40 | 82 | 0.4878 | 28.8821 | NS |
| IM62 | 5 | IM62+HWY | 20 | 54 | 108 | 162 | 0.6667 | 23.8263 | SIG |
| IM62 | 5 | IM62+HWY | 21 | 22 | 21 | 43 | 0.4884 | 28.3003 | NS |
| IM62 | 5 | IM62+HWY | 22 | 19 | 53 | 72 | 0.7361 | 29.2312 | SIG |
| IM62 | 4 | IM62+HWY | 23 | 30 | 86 | 116 | 0.7414 | 30.3648 | SIG |
| IM62 | 2 | IM62+HWY | 24 | 28 | 22 | 50 | 0.4400 | 28.9807 | NS |
| IM62 | 13 | IM62+HWY | 25 | 42 | 43 | 85 | 0.5059 | 30.1018 | NS |
| IM62 | 13 | IM62+HWY | 26 | 43 | 47 | 90 | 0.5222 | 32.1721 | NS |
| IM62 | 9 | IM62+HWY | 27 | 7 | 29 | 36 | 0.8056 | 24.5183 | NS |
| IM62 | 0 | IM62+HWY | 28 | 77 | 67 | 144 | 0.4653 | 22.7292 | NS |
| IM62 | 12 | IM62+HWY | 29 | 27 | 34 | 61 | 0.5574 | 21.2397 | NS |
| IM62 | 12 | IM62+HWY | 30 | 48 | 53 | 101 | 0.5248 | 24.7898 | NS |
| OD11 | 5 | IM62+OD | 1 | 1 | 27 | 28 | 0.9643 |  | SIG |
| OD11 | 2 | IM62+OD | 2 | 8 | 84 | 92 | 0.9130 |  | SIG |
| OD11 | 3 | IM62+OD | 3 | 13 | 87 | 100 | 0.8700 |  | NA |
| IM62 | 0 | IM62 | 1 | 164 | 8 | 172 | 0.0465 | 21.5154 | NA |
| IM62 | 7 | IM62 | 2 | 69 | 10 | 79 | 0.1266 | 26.9359 | NA |
| IM62 | 16 | IM62 | 3 | 154 | 8 | 162 | 0.0494 | 22.1432 | NA |
| IM62 | 0 | IM62 | 4 | 115 | 5 | 120 | 0.0417 | 25.9444 | NA |
| IM62 | 0 | IM62 | 5 | 24 | 9 | 33 | 0.2727 | 23.4533 | NA |
| IM62 | 7 | IM62 | 6 | 136 | 5 | 141 | 0.0355 | 17.7310 | NA |
| IM62 | 16 | IM62 | 7 | 140 | 4 | 144 | 0.0278 | 22.7858 | NA |
| IM62 | 11 | IM62 | 8 | 120 | 3 | 123 | 0.0244 | 26.9818 | NA |
| IM62 | 10 | IM62 | 9 | 101 | 3 | 104 | 0.0288 | 26.8373 | NA |
| IM62 | 13 | IM62 | 10 | 58 | 2 | 60 | 0.0333 | 32.8099 | NA |
| IM62 | 13 | IM62 | 11 | 51 | 5 | 56 | 0.0893 | 35.3827 | NA |
| IM62 | 11 | IM62 | 12 | 98 | 3 | 101 | 0.0297 | 22.9827 | NA |
| IM62 | 6 | IM62 | 13 | 96 | 4 | 100 | 0.0400 | 24.3869 | NA |
| IM62 | 6 | IM62 | 14 | 50 | 1 | 51 | 0.0196 | 27.9854 | NA |
| IM62 | 6 | IM62 | 15 | 109 | 0 | 109 | 0.0000 | 24.8258 | NA |
| IM62 | 6 | IM62 | 16 | 55 | 3 | 58 | 0.0517 | 27.3090 | NA |
| IM62 | 6 | IM62 | 17 | 83 | 0 | 83 | 0.0000 | 25.7117 | NA |
| IM62 | 4 | IM62 | 18 | 44 | 1 | 45 | 0.0222 | 27.6953 | NA |
| IM62 | 6 | IM62 | 19 | 90 | 3 | 93 | 0.0323 | 19.6464 | NA |
| IM62 | 13 | IM62 | 20 | 25 | 2 | 27 | 0.0741 | 28.5040 | NA |
| IM62 | 16 | IM62 | 21 | 112 | 14 | 126 | 0.1111 | 21.8118 | NA |
| IM62 | 7 | IM62 | 22 | 59 | 4 | 63 | 0.0635 | 20.1915 | NA |
| IM62 | 10 | IM62 | 23 | 74 | 2 | 76 | 0.0263 | 27.7943 | NA |
| IM62 | 10 | IM62 | 24 | 95 | 1 | 96 | 0.0104 | 22.5340 | NA |
| IM62 | 9 | IM62 | 25 | 92 | 0 | 92 | 0.0000 | 19.2019 | NA |
| IM62 | 9 | IM62 | 26 | 146 | 4 | 150 | 0.0267 | 25.5663 | NA |
| IM62 | 9 | IM62 | 27 | 84 | 5 | 89 | 0.0562 | 29.6586 | NA |
| IM62 | 16 | IM62 | 28 | 157 | 2 | 159 | 0.0126 | 25.3686 | NA |
| IM62 | 16 | IM62 | 29 | 26 | 6 | 32 | 0.1875 | 26.4208 | NA |
| IM62 | 16 | IM62 | 30 | 81 | 4 | 85 | 0.0471 | 22.0754 | NA |
| IM62 | 11 | IM62 | 31 | 131 | 8 | 139 | 0.0576 | 20.5657 | NA |
| HWY15D | 6 | HWY15 | 1 | 152 | 22 | 174 | 0.1264 | 13.8366 | NA |
| HWY15D | 6 | HWY15 | 2 | 214 | 17 | 231 | 0.0736 | 13.6000 | NA |
| HWY15D | 4 | HWY15 | 3 | 15 | 3 | 18 | 0.1667 | 20.6085 | NA |
| HWY15D | 5 | HWY15 | 4 | 268 | 29 | 297 | 0.0976 | 15.1823 | NA |
| HWY15D | 6 | HWY15 | 5 | 50 | 9 | 59 | 0.1525 | 18.2932 | NA |
| HWY15D | 6 | HWY15 | 6 | 42 | 8 | 50 | 0.1600 | 21.6989 | NA |
| HWY15D | 4 | HWY15 | 7 | 279 | 27 | 306 | 0.0882 | 16.3628 | NA |
| HWY15D | 4 | HWY15 | 8 | 186 | 24 | 210 | 0.1143 | 15.4090 | NA |
| HWY15D | 6 | HWY15 | 9 | 218 | 28 | 246 | 0.1138 | 15.2564 | NA |
| HWY15D | 6 | HWY15 | 10 | 180 | 21 | 201 | 0.1045 | 15.5260 | NA |
| HWY15D | 6 | HWY15 | 11 | 174 | 34 | 208 | 0.1635 | 13.7842 | NA |
| HWY15D | 7 | HWY15 | 12 | 153 | 18 | 171 | 0.1053 | 16.4726 | NA |
| HWY15D | 4 | HWY15 | 13 | 155 | 13 | 168 | 0.0774 | 17.1155 | NA |
| HWY15D | 4 | HWY15 | 14 | 208 | 21 | 229 | 0.0917 | 13.8453 | NA |
| HWY15D | 4 | HWY15 | 15 | 366 | 12 | 378 | 0.0317 | 14.6420 | NA |
| HWY15D | 4 | HWY15 | 16 | 261 | 32 | 293 | 0.1092 | 14.0407 | NA |
| HWY15D | 6 | HWY15 | 17 | 87 | 18 | 105 | 0.1714 | 14.8045 | NA |
| HWY15D | 4 | HWY15 | 18 | 349 | 27 | 376 | 0.0718 | 14.7766 | NA |
| HWY15D | 4 | HWY15 | 19 | 274 | 15 | 289 | 0.0519 | 16.2413 | NA |
| HWY15D | 6 | HWY15 | 20 | 222 | 34 | 256 | 0.1328 | 16.8549 | NA |
| HWY15D | 6 | HWY15 | 21 | 242 | 24 | 266 | 0.0902 | 15.4280 | NA |
| HWY15D | 6 | HWY15 | 22 | 49 | 9 | 58 | 0.1552 | 21.0781 | NA |
| HWY15D | 6 | HWY15 | 23 | 30 | 16 | 46 | 0.3478 | 20.7791 | NA |
| HWY15D | 6 | HWY15 | 24 | 322 | 9 | 331 | 0.0272 | 14.9536 | NA |
| OD11 | 3 | OD11 | 1 | 87 | 34 | 121 | 0.2810 | 11.4002 | NA |
| OD11 | 3 | OD11 | 2 | 104 | 32 | 136 | 0.2353 | 12.1900 | NA |
| OD11 | 5 | OD11 | 3 | 113 | 54 | 167 | 0.3234 | 16.5155 | NA |
| OD11 | 5 | OD11 | 4 | 137 | 44 | 181 | 0.2431 | 16.2901 | NA |
| OD11 | 5 | OD11 | 5 | 143 | 66 | 209 | 0.3158 | 15.7383 | NA |
| OD11 | 5 | OD11 | 6 | 16 | 8 | 24 | 0.3333 | 13.2572 | NA |
| OD11 | 3 | OD11 | 7 | 162 | 73 | 235 | 0.3106 | 13.5372 | NA |
| OD11 | 3 | OD11 | 8 | 15 | 5 | 20 | 0.2500 | 15.1527 | NA |
| OD11 | 3 | OD11 | 9 | 68 | 13 | 81 | 0.1605 | 13.1667 | NA |
| OD11 | 3 | OD11 | 10 | 50 | 16 | 66 | 0.2424 | 11.1004 | NA |
| OD11 | 3 | OD11 | 11 | 89 | 14 | 103 | 0.1359 | 12.2546 | NA |
| OD11 | 3 | OD11 | 12 | 88 | 18 | 106 | 0.1698 | 15.5921 | NA |
| OD11 | 5 | OD11 | 13 | 6 | 4 | 10 | 0.4000 | 14.5205 | NA |
| OD11 | 5 | OD11 | 14 | 78 | 43 | 121 | 0.3554 | 13.6518 | NA |
| OD11 | 2 | OD11 | 15 | 64 | 54 | 118 | 0.4576 | 15.6383 | NA |
| OD11 | 2 | OD11 | 16 | 76 | 34 | 110 | 0.3091 | 12.7179 | NA |
| OD11 | 2 | OD11 | 17 | 75 | 35 | 110 | 0.3182 | 13.2970 | NA |
| OD11 | 3 | OD11 | 18 | 104 | 52 | 156 | 0.3333 | 14.4804 | NA |
| OD11 | 3 | OD11 | 19 | 97 | 65 | 162 | 0.4012 | 12.8145 | NA |
| OD11 | 3 | OD11 | 20 | 35 | 21 | 56 | 0.3750 | 11.5924 | NA |

